# Association between genetically elevated levels of inflammatory biomarkers and risk of schizophrenia: a two-sample Mendelian randomisation study

**DOI:** 10.1101/123976

**Authors:** Fernando Pires Hartwig, Maria Carolina Borges, Bernardo Lessa Horta, Jack Bowden, George Davey Smith

**Author notes:** Corresponding author. Postgraduate Program in Epidemiology, Federal University of Pelotas, Pelotas (Brazil) 96020-220 and Medical Research Council Integrative Epidemiology Unit, University of Bristol, Bristol (United Kingdom) BS8 2BN.

## Abstract

**Background:** Positive associations between inflammatory biomarkers and risk of psychiatric disorders, including schizophrenia, have been reported in observational studies. However, conventional observational studies are prone to bias such as reverse causation and residual confounding.

**Methods:** In this study, we used summary data to evaluate the association of genetically elevated C reactive protein (CRP), interleukin-1 receptor antagonist (IL-1Ra) and soluble interleukin-6 receptor (IL-6R) levels with schizophrenia in a two-sample Mendelian randomisation design.

**Results:** The pooled odds ratio estimate using 18 CRP genetic instruments was 0.90 (95% CI: 0.84; 0.97) per two-fold increment in CRP levels; consistent results were obtained using different Mendelian randomisation methods and a more conservative set of instruments. The odds ratio for soluble IL-6R was 1.06 (95% CI: 1.01; 1.12) per two-fold increment. Estimates for IL-1Ra were inconsistent among instruments and pooled estimates were imprecise and centred on the null.

**Conclusion:** Under Mendelian randomisation assumptions, our findings suggest a protective causal effect of CRP and a risk-increasing causal effect of soluble IL-6R (potentially mediated at least in part by CRP) on schizophrenia risk.

## Introduction

Neuropsychiatric disorders are major contributors to the burden of disease worldwide due to its high impact on disability.^1,2^ Over one third of the top 25 global causes of disease burden in 2013, measured as years lived with disability (YLDs), falls into this category of disorders.^1^ Schizophrenia ranks 11^th^ among the leading global causes of YLDs^1^ and its high burden is associated with substantial personal and societal costs.^3^

A relation between schizophrenia and the immune system was suggested more than a century ago.^4^ Infections during pregnancy^5^ and early childhood,^6^ as well as autoimmune diseases,^7,8^ have been linked to increased risk of schizophrenia. In addition, findings from the largest genome-wide association study (GWAS) of schizophrenia to date corroborate that immune dysregulation plays a role in the pathogenesis of schizophrenia.^9^ Inflammation has been hypothesised as a potential mechanism linking the immune response to the pathogenesis of schizophrenia and other neuropsychiatric disorders since cytokines may influence multiple neurological processes, including neurotransmitter metabolism, neuroendocrine function, and neural plasticity.^4,10-12^ Observational epidemiological studies, mainly of cross-sectional design, indicate that circulating levels of several cytokines,^13^ such as interleukin (IL)-1β and IL-6, and of C reactive protein (CRP),^14,15^ are higher in schizophrenic individuals. Meta-analysis of randomised controlled trials suggested that anti-inflammatory drugs improve symptoms of the syndrome, but only a few studies with small sample sizes are available.^16^

Not only there are relatively few studies investigating the association between inflammatory biomarkers and schizophrenia in humans, but they are largely of observational nature. Conventional observational studies may have important limitations such as reverse causation and residual confounding,^17-19^ which hamper conclusions on whether specific anti-inflammatory agents could reduce risk of developing schizophrenia. Genetic variants can be used as instrumental variables of modifiable exposures in a Mendelian randomisation (MR) design to improve causal inference in observational studies. Justifications to rely on MR as a more robust method for causal inference than conventional observational studies include Mendel’s laws and the facts that genotypes of germline genetic variation are defined at conception and are generally not associated with conventional confounders of observational studies.^18-20^

MR has been used to investigate the causal effect of circulating CRP levels on schizophrenia risk. In a large Danish population-based study using four genetic instruments in the *CRP* gene region, the point estimate suggested a risk-increasing causal effect, but the lower limit of the 95% confidence intervals did not allow excluding the possibility of important protective effects.^21^ Two subsequent studies, both using the same summary association datasets in a two-sample MR design, reported directionally inconsistent estimates,^22,23^ likely due to a data harmonisation error in one of them.^24^ Moreover, neither of the two-sample MR studies performed substantial sensitivity analyses.

We investigated the causal effect of inflammatory markers on schizophrenia risk in a two-sample MR design.^25^

## Methods and Materials

### Datasets

We obtained summary association results for four sets of genetic instruments: CRP liberal (instruments selected using solely statistical criteria^26^), CRP conservative (instruments restricted to the *CRP* gene region^27^), Interleukin-1 receptor antagonist (IL-1Ra)^28^ and soluble Interleukin-6 receptor (sIL-6R).^29^ Summary associations between each instrument and schizophrenia risk were obtained from the largest schizophrenia GWAS to date^9^ (see Supplementary Methods for a description of each dataset). The summary genetic associations datasets were harmonised as described elsewhere,^24^ and are shown in Supplementary Tables 1 and 2.

**Table 1.** Odds ratio (OR) and 95% confidence intervals (95% CI) of schizophrenia per two-fold increments in inflammatory markers based on Mendelian randomisation (MR).

| SNP set | MR method | Parameter | OR or Odds<br>(95% CI) | P |
| --- | --- | --- | --- | --- |
| CRP liberal<br>(n=18)<sup>a</sup> | IVW (FE) | OR | 0.90 (0.86; 0.95) | 0.0004 |
|  | IVW (RE) | OR | 0.90 (0.84; 0.97) | 0.0054 |
|  | MR-Egger (FE) | Intercept (Odds) | 1.00 (0.99; 1.00) | 0.4245 |
|  |  | OR | 0.93 (0.85; 1.01) | 0.0900 |
|  | MR-Egger (RE) | Intercept (Odds) | 1.00 (0.99; 1.01) | 0.5657 |
|  |  | OR | 0.93 (0.82; 1.05) | 0.2149 |
|  | Weighted median | OR | 0.91 (0.85; 0.98) | 0.0159 |
| CRP conservative<br>(n=4)<sup>b</sup> | Ratio – rs1130864 | OR | 0.92 (0.81; 1.05) | 0.2112 |
|  | Ratio – rs1205 | OR | 0.92 (0.84; 1.01) | 0.0649 |
|  | Ratio – rs1800947 | OR | 0.90 (0.79; 1.02) | 0.0991 |
|  | Ratio – rs3093077 | OR | 0.97 (0.84; 1.12) | 0.6942 |
|  | WGLR | OR | 0.93 (0.86; 1.00) | 0.0418 |
| IL-1Ra (n=2) | Ratio – rs1542176 | OR | 1.02 (0.98; 1.07) | 0.3458 |
|  | Ratio – rs6743376 | OR | 0.98 (0.94; 1.01) | 0.2008 |
|  | IVW (FE) | OR | 0.99 (0.83; 1.19) | 0.7325 |
|  | IVW (RE) | OR | 0.99 (0.76; 1.30) | 0.8185 |
| IL-6R (n=1) | Ratio – rs222814 | OR | 1.06 (1.01; 1.12) | 0.0201 |

SNP: single nucleotide polymorphism. CRP: C reactive protein. IVW: inverse variance weighting. FE: fixed effects. RE: random effects. WGLR: Weighted generalised linear regression.
<sup>a</sup>The liberal CRP set corresponds to the SNPs identified by Dehghan and colleagues<sup>26</sup>.
<sup>b</sup>The conservative CRP set corresponds to the SNPs analysed by Wensley and colleagues<sup>27</sup>.

**Table 2.** Odds ratio (OR) and 95% confidence intervals (95% CI) of schizophrenia per two-fold increments in CRP levels based on Mendelian randomisation using the liberal set of 18 CRP-associated variants in a leave-one-out approach.

| Excluded<br>SNP | IVW (RE) |  | MR-Egger (RE) |  | Weighted median |  |
| --- | --- | --- | --- | --- | --- | --- |
|  | OR (95% CI) | *P* | OR (95% CI) | *P* | OR (95% CI) | *P* |
| rs10521222 | 0.90 (0.84; 0.97) | 0.007 | 0.93 (0.82; 1.05) | 0.228 | 0.91 (0.85; 0.97) | 0.015 |
| rs10745954 | 0.91 (0.85; 0.97) | 0.007 | 0.91 (0.81; 1.04) | 0.147 | 0.91 (0.86; 0.98) | 0.017 |
| rs1183910 | 0.91 (0.84; 0.98) | 0.019 | 0.94 (0.82; 1.08) | 0.365 | 0.94 (0.87; 1.01) | 0.099 |
| rs12037222 | 0.90 (0.84; 0.97) | 0.008 | 0.93 (0.81; 1.06) | 0.232 | 0.91 (0.86; 0.98) | 0.018 |
| rs12239046 | 0.90 (0.84; 0.97) | 0.007 | 0.93 (0.82; 1.06) | 0.253 | 0.91 (0.85; 0.97) | 0.015 |
| rs1260326 | 0.90 (0.84; 0.97) | 0.007 | 0.93 (0.82; 1.05) | 0.230 | 0.91 (0.85; 0.97) | 0.011 |
| rs13233571 | 0.90 (0.84; 0.96) | 0.005 | 0.93 (0.82; 1.05) | 0.242 | 0.91 (0.85; 0.97) | 0.016 |
| rs1800961 | 0.90 (0.84; 0.96) | 0.005 | 0.93 (0.82; 1.05) | 0.206 | 0.91 (0.85; 0.98) | 0.017 |
| rs2794520 | 0.90 (0.83; 0.98) | 0.014 | 0.93 (0.80; 1.08) | 0.304 | 0.94 (0.87; 1.01) | 0.128 |
| rs2847281 | 0.90 (0.84; 0.96) | 0.005 | 0.94 (0.83; 1.07) | 0.350 | 0.91 (0.85; 0.97) | 0.015 |
| rs340029 | 0.90 (0.84; 0.97) | 0.006 | 0.94 (0.82; 1.07) | 0.302 | 0.91 (0.85; 0.97) | 0.014 |
| rs4129267 | 0.91 (0.85; 0.98) | 0.011 | 0.93 (0.82; 1.05) | 0.226 | 0.92 (0.86; 0.99) | 0.030 |
| rs4420065 | 0.91 (0.84; 0.98) | 0.012 | 0.93 (0.82; 1.06) | 0.245 | 0.92 (0.86; 0.99) | 0.036 |
| rs4420638 | 0.88 (0.81; 0.95) | 0.002 | 0.87 (0.74; 1.02) | 0.082 | 0.89 (0.83; 0.95) | 0.004 |
| rs4705952 | 0.90 (0.84; 0.97) | 0.007 | 0.93 (0.82; 1.06) | 0.253 | 0.91 (0.85; 0.97) | 0.015 |
| rs6734238 | 0.90 (0.84; 0.97) | 0.007 | 0.93 (0.82; 1.06) | 0.252 | 0.91 (0.85; 0.97) | 0.013 |
| rs6901250 | 0.90 (0.84; 0.97) | 0.008 | 0.92 (0.81; 1.05) | 0.212 | 0.91 (0.85; 0.98) | 0.019 |
| rs9987289<sup>a</sup> | 0.91 (0.87; 0.96) | 0.002 | 0.92 (0.84; 1.01) | 0.093 | 0.91 (0.86; 0.98) | 0.018 |

SNP: single nucleotide polymorphism. CRP: C reactive protein. IVW: inverse variance weighting. RE: random effects.
<sup>a</sup>This variant was classified as potentially influential both in IVW and MR-Egger.

### Statistical analysis

SNP-biomarker associations were collected in ln-transformed units. Odds ratio estimates of schizophrenia per two-fold increments in circulating inflammatory biomarker levels were obtained as follows: 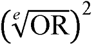, where OR is the odds ratio estimate per 1-ln increment in biomarker levels, and *e* is the base of the natural logarithm.

MR requires that a) the genetic instruments are associated with the modifiable exposure of interest, and b) any association between the instruments and the outcome is entirely mediated by the exposure. Given that assumption a) is empirically verifiable, careful consideration of potential violations of assumption b) (due to factors such as population stratification, linkage disequilibrium, canalisation or horizontal pleiotropy) is important to minimise bias.^18^

Five MR methods were used:

i) *Ratio method*: this method was used to obtain individual-SNP estimates by dividing the SNP-schizophrenia by the corresponding SNP-biomarker effect estimates. Standard errors were estimated using the Delta method^30^ assuming the uncertainty in the SNP-exposure association estimates was negligible (the NO Measurement Error – NOME – assumption). These standard errors were then used to perform weighted analyses using methods (ii)-(iv).
ii) *Inverse variance weighting* (*IVW*): IVW is equivalent to the inverse variance weighted average of ratio estimates from two or more instruments. It was implemented as a weighted linear regression of SNP-schizophrenia against SNP-biomarker effect estimates with the intercept constrained at zero, which follows from the assumption that there is no overall unbalanced horizontal pleiotropy.^25^ We performed both fixed and multiplicative random effects IVW, since the fixed effects method may be over-precise in the presence of heterogeneity (which can occur due to, among other factors, horizontal pleiotropy).^31^
iii) *Weighted generalised linear regression* (*WGLR*): this method is similar to the IVW method, but allows properly accounting for the correlation between the genetic instruments.^32^ The WLGR method was used instead of the IVW method when using the conservative set of CRP genetic instruments, which comprised variants in partial linkage.
iv) *Weighted median*: The weighted median estimate is the median of the weighted empirical distribution function of individual-SNP ratio estimates. It differs from a simple median estimate because the amount of information that a given instrument contributes to the estimate depends on the precision of its ratio estimate. This method provides a consistent causal effect estimate if more than 50% of the information comes from valid SNPs.^33^
v) *MR-Egger regression*: MR-Egger implementation consists of the same weighted linear regression model used in IVW, but the intercept is not constrained. Assuming that horizontal pleiotropic effects and SNP-exposure associations are independent (i.e., the InSIDE assumption), MR-Egger regression provides a valid causal effect estimate even if all SNPs are consistent. Moreover, its intercept can be interpreted as a test of overall unbalanced horizontal pleiotropy.^34^ Both fixed and multiplicative random effects versions of the MR-Egger regression method were performed.

Measurement error in the SNP-exposure associations is always present to some degree. In the two-sample setting, it attenuates the causal effect estimates, and also affects MR-Egger regression intercept. The degree of NOME violation in IVW and MR-Egger regression can be quantified by the 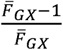 and 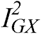 statistics, respectively.^35^ Both range from 0% to 100%, and can be interpreted as the amount of attenuation in the effect estimates due to NOME violations.^35,36^ Such violations can be accounted for using the Simulation Extrapolation (SIMEX) method, which was applied to MR-Egger regression.^36^

The Cochran’s Q test for heterogeneity was applied to the CRP liberal set to test for the presence of horizontal pleiotropy. This test assumes that all valid genetic instruments estimate the same causal effect.^37^ Moreover, to identify potentially influential instruments in the statistical set of CRP instruments, two tests of influence (based on studentised residuals or Cook’s distance) were applied separately for IVW or MR-Egger regression.^38,39^ P-values for the studentised residuals test were obtained from a Student’s t-distribution with degrees of freedom equal to *L*−2 (for IVW) or *L*−3 (for MR-Egger regression), with *L* being the number of genetic instruments. The F distribution with joint degrees of freedom equal to (1, *L*−1) (for IVW) or (1, *L*−2) (for MR-Egger regression) was used for the Cook’s distance test. We then applied three different statistical significance criteria to classify SNPs as potentially influent: *P*<0.01, *P*<0.05 n or *P*<0.1 in at least one of the influence tests. A leave-one-out approach to evaluate the influence of each instrument was also performed.

We used available data on the association between the sIL-6R genetic instrument and CRP levels^29^ in a mediation analysis evaluating the potential mediating effect of CRP in the association between sIL-6R levels and schizophrenia risk. We used MR to obtain causal effect estimates of the exposure-outcome (i.e., sIL-6R levels and schizophrenia risk), exposure-mediator (i.e., sIL-6R levels and CRP levels) and mediator-outcome (i.e., CRP levels and schizophrenia risk) associations. The last two can be used to estimate the expected effect of sIL-6R levels on schizophrenia risk assuming that CRP levels fully mediates this association. This can then be contrasted to the observed exposure-outcome association.^40^ More specifically, the expected effect assuming full mediation (*β*_*E*_) can be estimated as 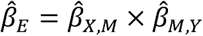, where 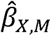 is the causal effect estimate of the exposure on the mediator, and 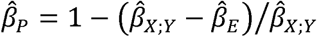 is the causal effect estimate of the mediator on the outcome. The proportion of the effect of the exposure on the outcome that is mediated by the mediator (*β*_*P*_) can then be estimated as 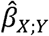, where 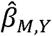 is the causal effect estimate of the exposure on the outcome. Standard errors for 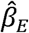 and 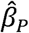 were estimated using parametric bootstrap, and confidence intervals were derived using the normal approximation method.

Analyses were performed using R 3.2.4 (www.r-project.org).

## Results

Table 1 displays the association between genetically elevated inflammatory biomarkers and schizophrenia. Regarding CRP, when the liberal set of 18 CRP-associated variants was used, results were consistent among the three MR methods, with odds ratio of schizophrenia of 0.90 (random effects 95% CI: 0.84; 0.97), 0.91 (95% CI: 0.85; 0.98) and 0.93 (random effects 95% CI: 0.82; 1.05) per two-fold increment in circulating CRP levels using IVW, weighted median and MR-Egger regression approaches, respectively (Figure 1). The 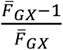 and 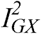 statistics were 99.8% and 98.3%, respectively, suggesting that measurement error in the SNP-CRP associations were not substantially attenuating the causal effect estimates. Indeed, regular and SIMEX-corrected MR-Egger regression results were virtually identical, so only the first was shown.

**Figure 1.**
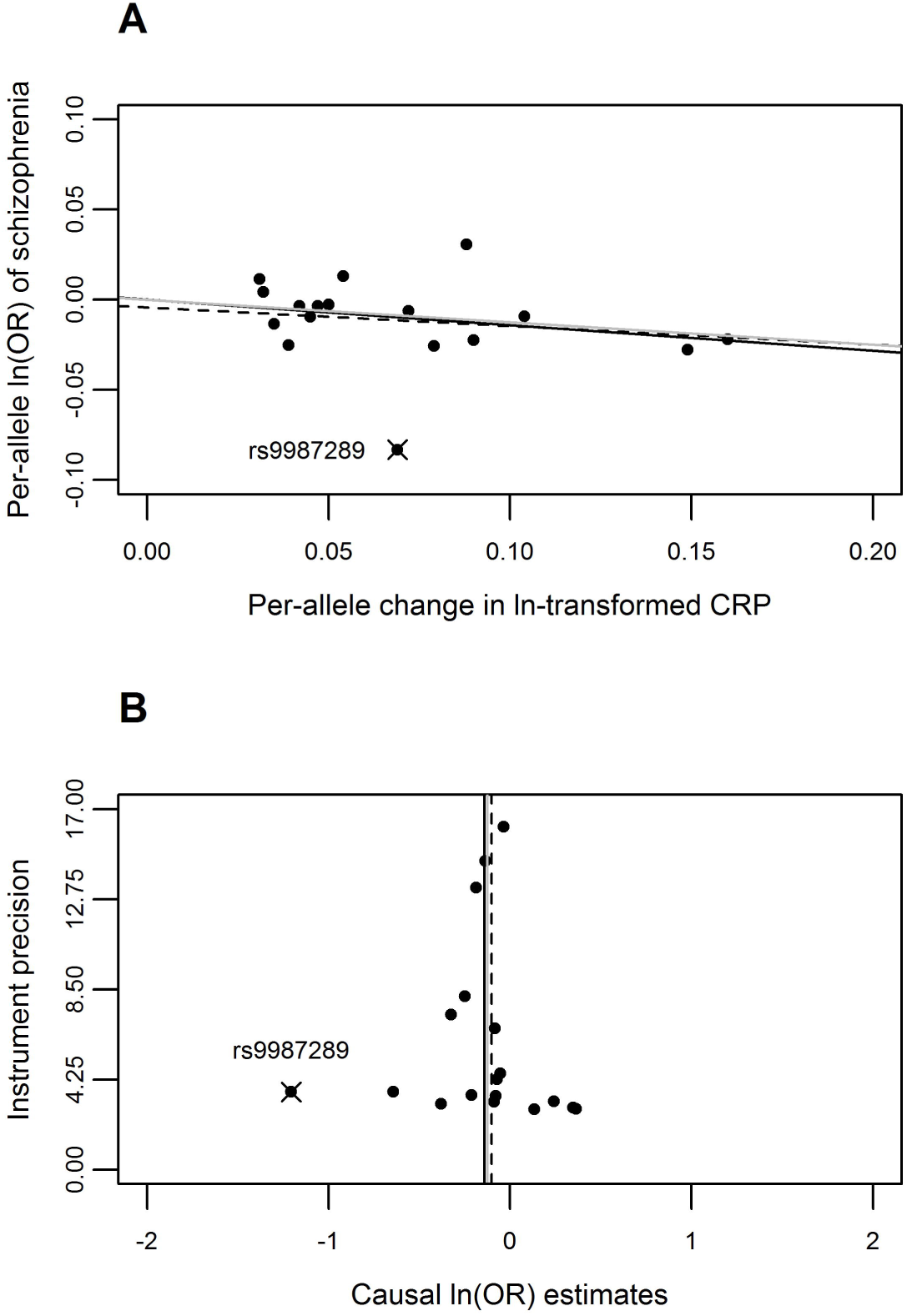
Scatter and funnel plots of the Mendelian randomisation analyses regarding the causal effect of C reactive protein levels (CRP) on schizophrenia using the liberal set of 18 genetic instruments. IVW, MR-Egger and weighted median estimates are indicated in solid, dashed and grey lines, respectively. The single genetic instrument classified as influential (rs9987289) was marked with an “X” and labeled. A: scatter plot of instrument-schizophrenia associations in the y-axis against instrument-CRP associations in the x-axis. B: funnel plot of instrument precision (i.e., instrument-CRP regression coefficients divided by the correspondent instrument-schizophrenia standard errors) in the y-axis against individual-instrument ratio estimates in log odds ratio of schizophrenia in the x-axis.

MR-Egger intercepts were (when rounding to two decimal digits) equal to 1.00 with narrow confidence intervals, suggesting no strong overall unbalanced horizontal pleiotropy. However, the Cochran’s Q statistics was 31.9, with an associated P-value of 0.015, suggesting some heterogeneity in the causal effect estimates, possibly due to horizontal pleiotropy.

The results obtained using the conservative CRP set were consistent with the liberal set. Odds ratio estimates of schizophrenia per two-fold increment in circulating CRP levels based on the ratio method ranged from 0.90 (95% CI: 0.79; 1.02) to 0.97 (95% CI: 0.84; 1.12). The pooled estimate was 0.93 (95% CI: 0.82; 1.05). The two estimates for IL-1Ra were directionally inconsistent (1.02 and 0.98), with a pooled IVW estimate of 0.99 (random effects 95% CI: 0.76; 1.30). Finally, odds ratio of schizophrenia per two-fold increments in circulating sIL-6R levels were 1.06 (95% CI: 1.01; 1.12).

In the leave-one-out MR analyses using the liberal CRP set, all odds ratio estimates of schizophrenia per two-fold increment in circulating CRP levels were directionally consistent (Table 2). IVW, weighted median, and MR-Egger regression estimates ranged from 0.88 to 0.91, 0.89 to 0.94 and 0.87 to 0.94, respectively. Conventional statistical significance levels were achieved in all IVW estimates and in 16 weighted median estimates, but in none of the MR-Egger regression estimates. The SNP rs9987289 was the only variant classified as potential influential, and its removal had virtually no effect on the results. This single variant accounted for most of the heterogeneity in the individual-instrument ratio estimates (Supplementary Table 4). Therefore, any possible horizontal pleiotropy suggested by the Cochran’s Q test does not explain our findings.

Using the ratio method, the causal effect of 1-unit increase in ln(sIL-6R) levels on ln(CRP) levels was -0.26 (95% CI: -0.32; -0.21), and the causal effect of 1-unit increase in ln(CRP) levels on ln(odds ratio) of schizophrenia was -0.14 (95% CI: -0.23; -0.05). Multiplying these estimates yielded an odds ratio of schizophrenia of 1.03 (95% CI: 1.01; 1.04) per two-fold increment in sIL-6R levels. The estimated proportion of the effect of sIL-6R levels on schizophrenia risk that is mediated by CRP (using the IVW estimate from the liberal CRP set as the causal effect estimate of CRP on schizophrenia risk) was 43.8% (95% CI: 3.3%; 84.2%).

## Discussion

We used two-sample MR to evaluate the association of inflammatory biomarkers with schizophrenia. Overall, we did not find strong evidence that lifelong exposure to increased action of these pro-inflammatory cytokines increases schizophrenia risk, as previously hypothesised, and, conversely, found that blockade of IL-6 effects and low CRP levels might in fact increase schizophrenia risk. Moreover, part of the relation between IL-6 signalling and schizophrenia might be mediated by CRP levels, which is consistent with previous knowledge on the major role of IL-6 in inducing acute phase response and the fact that lower CRP levels are a downstream effect of inhibiting IL-6 classical signalling.^26,41,42^ For IL-1Ra, point estimates were inconsistent between instruments and confidence intervals were large. Our study extends previous MR findings by evaluating different inflammatory markers and applying a range of sensitivity analyses.

IL-6 is known for its pivotal role in integrating immune response, such as by inducing hepatic acute-phase proteins, differentiation of T-cells, and tissue regeneration.^43^ Apart from being a sensitive marker of systemic inflammation and tissue damage, CRP is an acute phase protein that contributes to host defence against infection.^44^ CRP binds to phosphocholine expressed on the surface of cells and some bacteria, which activates the complement system, promoting phagocytosis and clearance of necrotic/apoptotic cells and bacteria.^44,45^

Mechanisms underlying the association of blockade of IL-6 classical signalling and lower CRP levels with increased risk of schizophrenia are unknown. We speculate that they relate to increased susceptibility to early life infection. Blockade of IL-6 classic signalling leads to increased susceptibility to infections in mice^46^ and humans.^43^ Observational studies indicate that low levels of some acute phase proteins in newborns were related to higher schizophrenia risk,^47^ and that neonates that develop schizophrenia later in life have an impaired capacity of increasing levels of acute phase proteins, such as CRP, in response to some maternal infections compared to controls.^48^ In adults, prospective studies indicate that higher CRP levels are related to increased susceptibility to infections.^49,50^ However, these findings should be interpreted cautiously as higher CRP may simply reflect subclinical infection, chronic activation of the inflammatory response, pre-existing disease, socio-economic or lifestyle characteristics. A MR study reported that genetic predisposition to higher CRP levels was not associated with increased infection risk in adults^50^ in adults. To the best of our knowledge, no existing MR study has investigated IL-6 and CRP effects on early life infection risk.

Causal inference from MR relies on some assumptions, one of them being that the exposure completely mediates the association (if any) between the instrument(s) and the outcome. Most of the genetic instruments that we used have biological justifications for their selection, except for the liberal CRP set of 18 variants. Indeed, half of the variants in the liberal CRP set have been associated with one or more of 10 tested cardiometabolic phenotypes, while the other half was not. Among the first half of variants, six (or other variants indexing the same locus) were associated with CRP levels independently on their association with cardiometabolic phenotypes, which was not the case for the remaining three variants.^42^ Although these findings suggest that the CRP-associated SNPs in the liberal set are pleiotropic, our results based on these variants were consistent among the three MR methods (which have different assumptions regarding horizontal pleiotropy) and were corroborated by the results for the conservative CRP set and the leave-one-out analysis. The latter is also important because the liberal CRP set included the IL-6R variant rs4129267, which was not used as a genetic instrument for IL-6R, but might nevertheless influence sIL-6R levels. Moreover, previous studies using the conservative CRP set, the IL-R1a and the IL-6R instruments observed that, in general, those variants are not associated with conventional confounders. ^27,29,51^

SNP-biomarker and SNP-schizophrenia estimates were obtained in mostly European studies, thus minimizing the possibility population stratification bias. This also increases the plausibility of the two-sample MR assumption that summary genetic association results were obtained in samples from the same or comparable populations. Regarding power, although some SNP-biomarker associations were estimated in small samples, the statistical evidence for association in such datasets was generally strong. Moreover, power in the two-sample setting depends more on the precision of the SNP-outcome than on the SNP-exposure association,^25^ and SNP-schizophrenia associations were estimated in ∼80,000 individuals.

Interpreting the magnitude of estimates for the effect of CRP and IL-6 on schizophrenia risk, as well as for the mediated effect of IL-6 by circulating CRP, requires caution. Our MR analysis likely reflects lifelong exposure to elevated cytokines and CRP. However, it is possible that only exposure to IL-6 and CRP in a specific window of time (e.g., early life) affects schizophrenia risk. We obtained estimates for the SNP-cytokines and SNP-CRP associations from adult subjects, but these associations might differ in early life. In addition, we used estimates for the effect of the IL-6R genetic instrument on sIL- 6R to investigate the total and the indirect (mediated by CRP) effect of blocking IL-6 classical signalling. However, this genetic instrument affects IL-6 classical signalling by increasing cleavage of membrane-bound IL-6R, which results in lower availability of membrane-bound IL-6R and higher availability of soluble IL-6R. Both mechanisms are likely to contribute to inhibiting IL-6 classical signalling.^43^ Finally, it is possible that IL- 6 and CRP effects on schizophrenia risk are related to a maternal effect (e.g., maternal susceptibility to infections during pregnancy), so that our findings are explained by the correlation between maternal and offspring genotypes.

Our findings support the notion that lower CRP levels and blockade of IL-6 cell signalling, both associated with lower inflammation and acute phase response, increase schizophrenia risk. This suggests that the positive associations of CRP and IL-6 with schizophrenia risk in conventional observational studies are due to limitations such as reverse causation or residual confounding. Even though our findings could be a result of horizontal pleiotropy that we failed to detect and account for, they at least suggest that increased levels of inflammatory biomarkers do not lead to substantially higher schizophrenia risk.

## Acknowledgements

The Medical Research Council (MRC) and the University of Bristol fund the MRC Integrative Epidemiology Unit [MC_UU_12013/1, MC_UU_12013/9]. The funders played no role in the design, analysis or interpretation of the findings from this study.

